# Resting-state functional connectivity changes following audio-tactile speech training

**DOI:** 10.1101/2024.10.26.620393

**Authors:** Katarzyna Cieśla, Tomasz Wolak, Amir Amedi

## Abstract

Understanding speech in background noise is a challenging task, especially if the signal is also distorted. In a series of previous studies we have shown that comprehension can improve if simultaneously to the auditory speech, the person receives speech-extracted low-frequency signals on fingertips. The effect increases after short audio-tactile speech training. Here we use resting-state functional magnetic resonance, measuring spontaneous low-frequency oscillations in the brain while at rest, to assess training-induced changes in functional connectivity. We show enhanced connectivity within a right-hemisphere cluster encompassing the middle temporal motion area (MT), and the extrastriate body area (EBA), and lateral occipital cortex (LOC), which before training is found to be more connected to bilateral dorsal anterior insula. Furthermore, early visual areas are found to switch from increased connectivity with the auditory cortex before, to increased connectivity with an association sensory/multisensory parietal hub, contralateral to the palm receiving vibrotactile inputs, after. Also the right sensorimotor cortex, including finger representations, is more connected internally after training. The results alltogether can be interpreted within two main complementary frameworks. One, speech-specific, relates to the pre-existing brain connectivity for audio-visual speech processing, including early visual, motion and body regions for lip-reading and gesture analysis in difficult acoustic conditions, which the new audio-tactile speech network might be built upon. The other refers to spatial/body awareness and audio-tactile integration, including in the revealed parietal and insular regions. It is possible that an extended training period may be necessary to more effectively strengthen direct connections between the auditory and sensorimotor brain regions, for the utterly novel speech comprehension task. The outcomes of the study can be relevant for both basic neuroscience, as well as development of rehabilitation tools for the hearing impaired population.

## Introduction

Resting-state functional magnetic resonance imaging (rsfMRI) is based on the premise that spontaneous low-frequency oscillations in the brain while at rest can reveal its functional organization and its alterations, without the need for the subject to engage in a specific task (Biswal et al., 1995; Raichle 2009). At the same time, the signals obtained from task-MRI and rsfMRI are likely to stem from similar structural connections and neuronal processes (Ngo et al. 2022). In addition, in resting-state protocols, the oscillations of the BOLD signal offer up to three times higher signal-to-noise ratio compared to the signal increases observed during task-related activities (Fox and Greicius, 2010). Using a measure of functional connectivity (FC; representing signal correlations among remote brain regions), rsMRI has been shown to reveal language networks with high sensitivity (eg. Lemme et al. 2019, Tie et al., 2014; Zhu et al., 2014, Branco et al. 2016, Sun et al. 2019), and considerable FC changes were reported as a result of e.g. auditory or language training, among other types of cognitive interventions (Fox and Greicius, 2010, Liu et al. 2021; Katsuno et al. 2022; Maruyama et al. 2018, Cao et al. 2016, Langer et al. 2013, Bamidis et al. 2014). It has been proposed that to gain a deeper understanding of improved performance following treatment, it is essential to consider functional brain networks, which represent a more complex level of brain organization.

In the current study we use the rsfMRI method specifically to investigate changes in functional connectivity following speech comprehension training, combining convergent auditory and vibrotactile inputs on fingertips.

Audio-(vibro)tactile interactions are relatively common in our everyday life, e.g. when receiving tactile feedback from a ringing phone, steering wheel, or when playing computer games. The combination and integration of these two sensory signals seems natural and inherent (van Bekesy 1959). This possibly due the vast similarities between the two sensory modalities, including the capability to encode the very same oscillatory patterns within an overlapping frequency range using mechanoreceptors (specifically for the tactile sense, Pacinian corpuscles on the skin respond to vibrations up to 700-1000 Hz, while the auditory system encodes vibrations in the air using inner hair cells between 20 Hz and 20 kHz; Bolanowski 1998, Schnupp, Nelken & King, 2011). Also evolutionary similarities exist between the two senses, with the organ of Corti in the inner ear speculated to have originally developed from epithelium (Pisciottano et al. 2019).

In several published works, convergent vibrotactile inputs have been shown to successfully enhance music perception (e.g. Russo, Ammirante & Fels, 2012; Sharp et al., 2019), as well as localization of sounds in space (Gori et al., 2014; Occelli, Spence & Zampini, 2011, Snir & Ciesla, et al. 2024). Another specific use of low-frequency vibrations has been to improve speech comprehension. This approach was originally developed to assist the deaf population, with several tactile aids designed to convey different features of speech signals via the skin (e.g. Galvin et al. 1991, Weisenberger et al. 1995). In a series of recent studies, by our lab and others, it was shown that improvement of speech perception is also possible in typically hearing and hearing impaired people using cochlear implants, with speech-extracted vibrations delivered on fingertips (Cieśla et al., 2019; Cieśla et al., 2022; Fletcher et al., 2019; Huang et al., 2017; Schulte et al. 2023, Rautu et al. 2023). Most authors implemented some form of manipulation to the speech signal to mimic the acoustic conditions that are most challenging for the hearing-impaired population, such as speech distortions and/or simultaneous background noise.

This rsfMRI study is complementary to our parallel task-fMRI study where participants were performing speech comprehension tasks inside an MRI scanner before and after dedicated audio-tactile speech comprehension training (Ciesla et al. 2024). The speech signals were distorted (vocoded to resemble speech delivered through a cochlear implant system) sentences presented within speech background noise. The tactile vibrations corresponded to low frequencies (<200Hz) extracted directly from the speech inputs. We demonstrated enhanced comprehension following audio-tactile training, which aligns with findings from several other studies that utilized audio-visual speech comprehension training, as well as those showing improved intelligibility of distorted speech with practice time (e.g. Casserly and Pisoni, 2015, Erb et al., 2013; Guediche et al. 2014, Bernstein 2013). The results suggest that the speech-extracted tactile signal can potentially work in a manner similar to the supporting visual signal (for example through lip-reading), through temporal synchronization with the auditory inputs (O’Sullivan 2021; Schulte et al. 2023).

Here we speculated that training-induced neuronal changes will also manifest in resting-state functional connectivity (FC). Specifically, we hypothesized to see enhanced connectivity between the auditory and the somatosensory systems, and possibly with other language-related areas (frontal, temporal, parietal; Price et al. 2012, Hertrich et al. 2020), reflecting familiarization with the new multisensory context of speech perception and perceptual learning. Since improvement in speech comprehension (as demonstrated in behavioral results, Ciesla et al. 2024) required successful multisensory integration, we furthermore expected to see changes in functional connectivity of multisensory regions, such as the posterior superior temporal sulcus (pSTS) or angular gyrus/supramarginal gyrus (AG/SMG), shown in the literature for both audio-visual and audio-tactile integration (e.g., Renier et al. 2009, Kassuba et al., 2013, Landelle et al. 2023; King et al. 2019; Nath et al. 2012; Hickock et al. 2018). The applied outcome measure was the difference in functional connectivity patterns between two resting-state fMRI sessions, before and after training.

## Material and Methods

The study took place at the ELSC Neuroimaging Unit in Jerusalem, Israel. All participants gave informed consent and received compensation for their participation. The study procedures complied with the Declaration of Helsinki (2013) and received approval from the Ethics Committee of the Hadassah Medical Center in Jerusalem (protocol no. 353-18.1.08).

## Participants

The reported data was acquired in seventeen adult healthy individuals (6 male/11 female, 26.8+/-4.2), all right-handed and with no history of neurological or neurodevelopmental impairments.

## Experimental procedures

### Behavioral experimental procedures

As already described in our parallel paper in detail (Ciesla et al. 2024), between the two resting-state fMRI sessions whose results we report here, the participants were tested twice (before and after training) in their vocoded speech-in-noise comprehension. There were three test conditions : a) auditory sentences (A), b) auditory sentences with congruent tactile vibrations delivered on two fingertips of the right hand (ATc), b) auditory sentences accompanied with non-congruent tactile vibrations on the fingertips (ATnc). For each test condition, an individual Speech Reception Threshold (SRT, i.e. SNR between the target sentence and the background noise, for 50% understanding) was determined. The applied sentences were derived from the HINT database (Nilsson et al. 1994) and vocoded to resemble stimulation through a cochlear-implant (Walkowiak et al. 2010). In the multisensory test conditions, low-frequency vibrations (<f0 of the spoken sentence, i.e. 200Hz) were extracted from either the concurrently presented auditory sentence (audio-tactile congruent, ATc), or from another randomly selected sentence from the HINT database (audio-tactile non-congruent, ATnc), and delivered on the index and middle finger of the right hand through an in-house developed device (Ciesla et al. 2022). Between the two SRT testing sessions, the participants took part in a 30–45 min training session. They were asked to repeat 148 HINT sentences, one by one, with accompanying congruent tactile vibrations delivered on the fingertips, at individually determined SRT. The training session continued until all 148 sentences were repeated correctly without feedback.

### Resting state fMRI data acquisition

Resting state fMRI data was aquired twice using a 3T Skyra MRI scanner by Siemens and a 32-channel head-coil, before and after training, during a 10:11 min session. The parameters were the following : TR =1.5s, TE=32.6ms, voxel size 2×2×2 mm, 72 slices, flip angle 78, FOV 192 mm, bandwidth 1736 Hz. Participants had their eyes open, and were asked to fixate on a cross presented on the screen. In addition, a high-resolution T1 MR sequence was collected with the following parameters: TR=2.3 s, TE=2.98 ms, IT=900ms, matrix size 256×256, voxel size : 1×1×1 mm, 160 slices, Bandwidth 240Hz/voxel.

## Data analysis

### Behavioral data analysis

The SRT values from participants’ speech comprehension tests before and after training were compared across different task conditions within the Pre or Post test sessions (A, ATc, ATnc; using Wilcoxon ranked-sum tests), as well as between sessions (A Pre vs A Post, ATc Pre vs ATc Post, ATnc Pre vs ATnc Post; using Wilcoxon signed-rank tests).

### Resting-state fMRI data analysis

In order to evaluate functional connectivity patterns between brain regions, CONN 22a (Connectivity Toolbox, https://web.conn-toolbox.org/) software was used for data analysis. Preprocessing of the functional data included: slice timing correction, motion correction, scrubbing, linear detrending, band-pass filtering (0.008 Hz < f < 0.09 Hz), co-registration to individual T1 structural scans, surface-based spatial normalization (https://surfer.nmr.mgh.harvard.edu/), and spatial smoothing (6 mm Gaussian kernel). Denoising was applied with default settings. A group-ICA analysis was performed using 40 components and the G1 FastICA + GICA3 back-projection procedure with 64 dimensionality reduction. Next, independent components for further analysis were selected using visual inspection (choosing those representing known networks or functional hubs, such as auditory, language, visual, sensorimotor, etc.). Functional connectivity maps were compared between sessions (PRE vs POST). The subsequent statistical analysis used permutation with the following parameters : voxel-threshold p <0.01, cluster-mass p-FDR corrected p < 0.05. The regions were labeled using the Harvard-Oxford Brain Atlas.

## Results

### Lowest SRT values show best performance in the congruent audio-tactile task after training

The participants were found to significantly improve in speech comprehension after training in both the auditory (A; by mean 10 dB) and audio-tactile congruent (ATc, by mean 8.5 dB) test conditions. In addition, lowest SRT values, indicating the same level of understanding (50%) in more difficult acoustic conditions, were found for the ATc test after training (2.6dB+/-6.3 dB). Statistical details can be found in (Ciesla et al. 2024).

### Resting-state fMRI functional connectivity changes after training in the auditory, sensorimotor and visual systems

We show that resting-state functional connectivity *increased* after, as compared to before, audio-tactile speech comprehension training between bilateral clusters encompassing parts of lateral occipital cortex (LOC)/angular gyrus (AG)/posterior medial temporal gyrus (pMTG)/posterior superior temporal sulcus (pSTS) (seed; including regions such as MT +/-44 -68 2, EBA -/+ 50 -70 5; https://neurosynth.org/), and LOC/pMTG/AG on the right side (target), indicating increased internal connectivity in this network; between bilateral clusters encompassing parts of the occipital pole (OP)/LOC/lingual gyrus (LG)/intracalcarine cortex (ICC) (early visual cortex; seed), and a hub representing left superior parietal lobe (SPL)/anterior supramarginal gyrus (aSMG) / postcentral gyrus (PostCG) (target); and of a right-hemisphere cluster encompassing parts of PostCG/precentral gurus (PreCG)/SPL (sensorimotor cortex) with itself. At the same time, *decreased* functional connectivity after training was revealed between bilateral clusters encompassing parts of iLOC/AG/pMTG/pSTS (same seed as above) and bilateral anterior insula (target), as well as between bilateral clusters encompassing parts of superior temporal gyrus (STG)/temporal pole (PT) /polar pole (PP)/SMG (auditory cortex), and OP/LOC on the right side (early visual cortex). Results are depicted in Figures 1-2.

**Figure 1.**
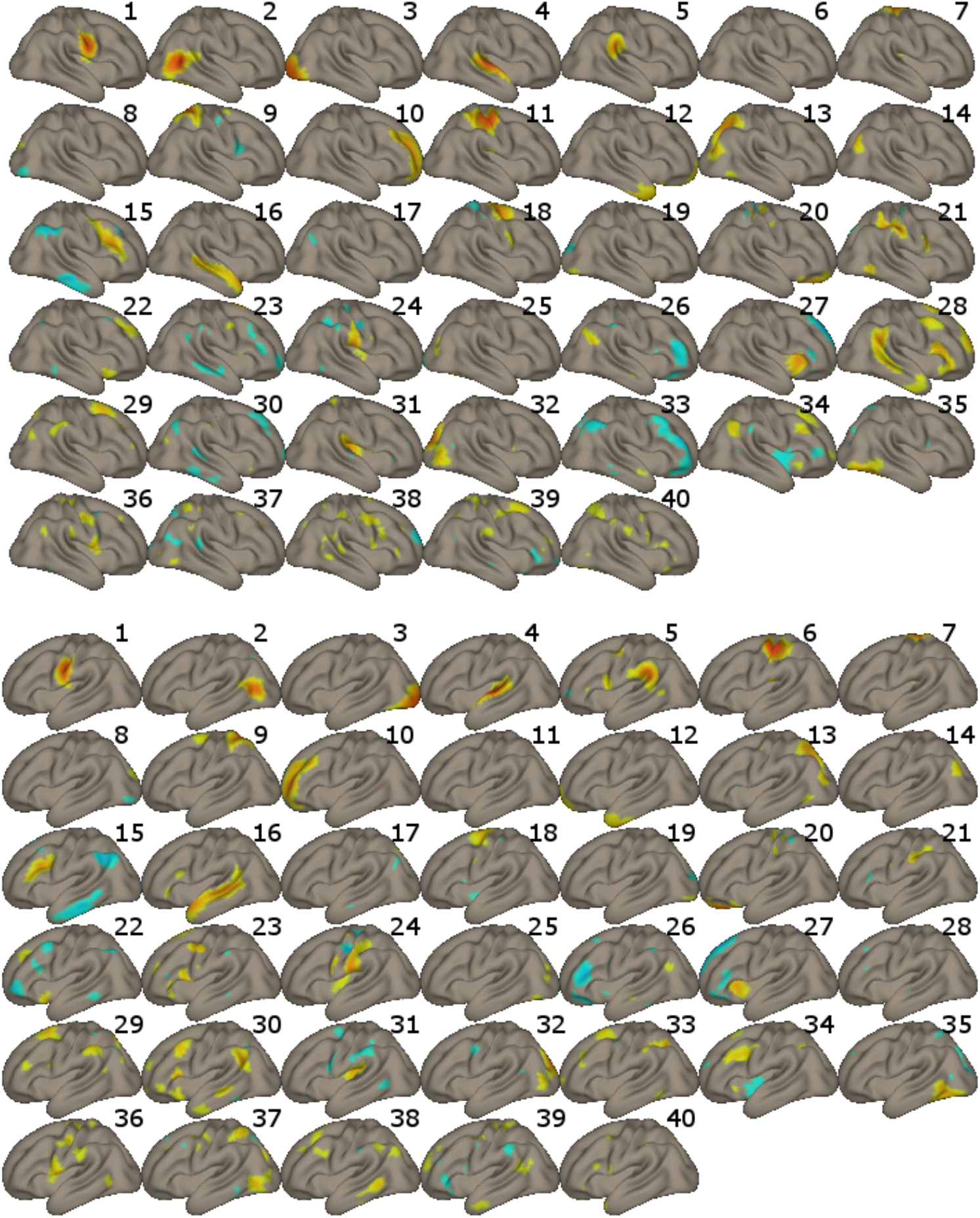
Forty Independent Components (ICs) revealed in the ICA analysis.

**Figure 2.**
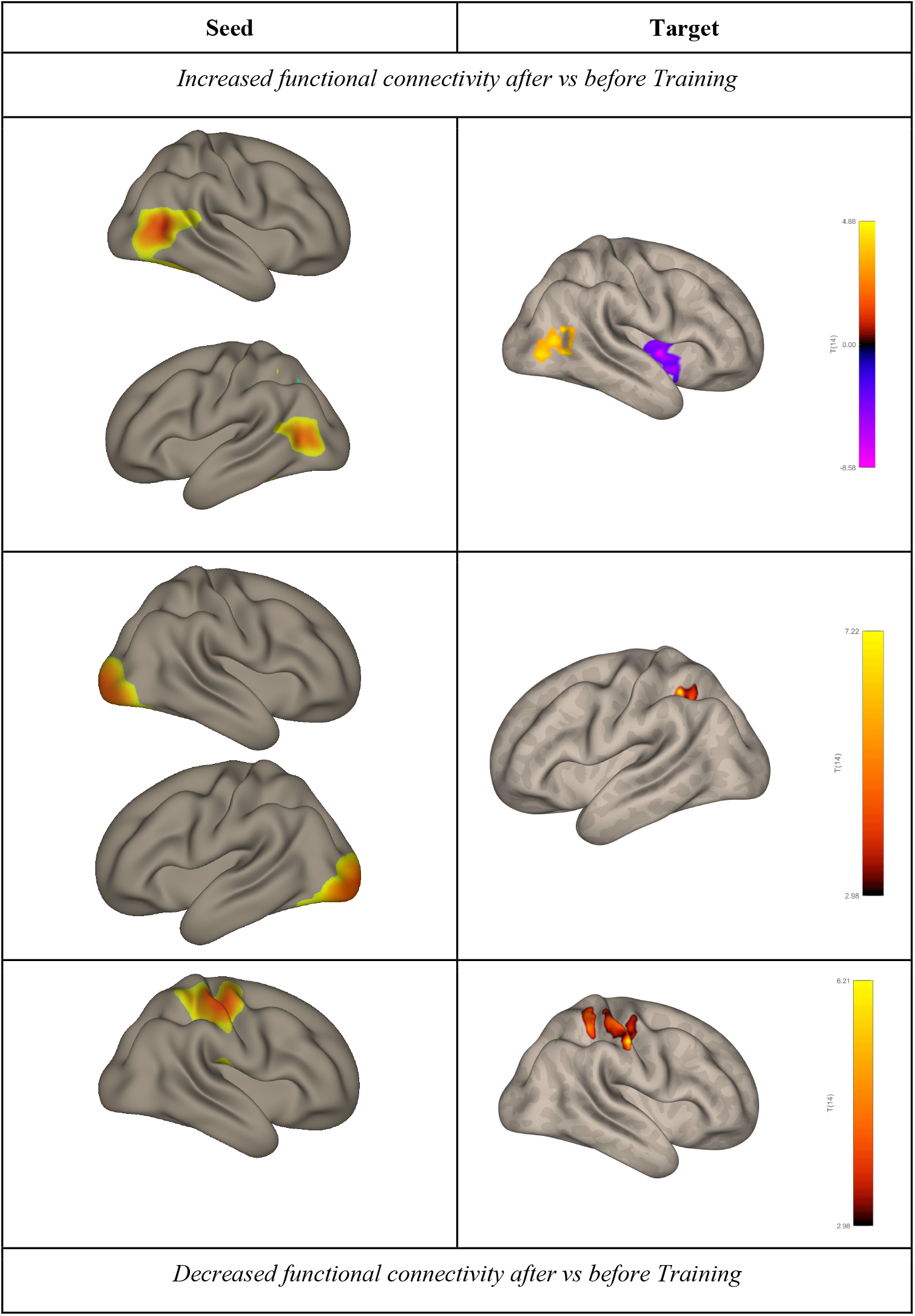
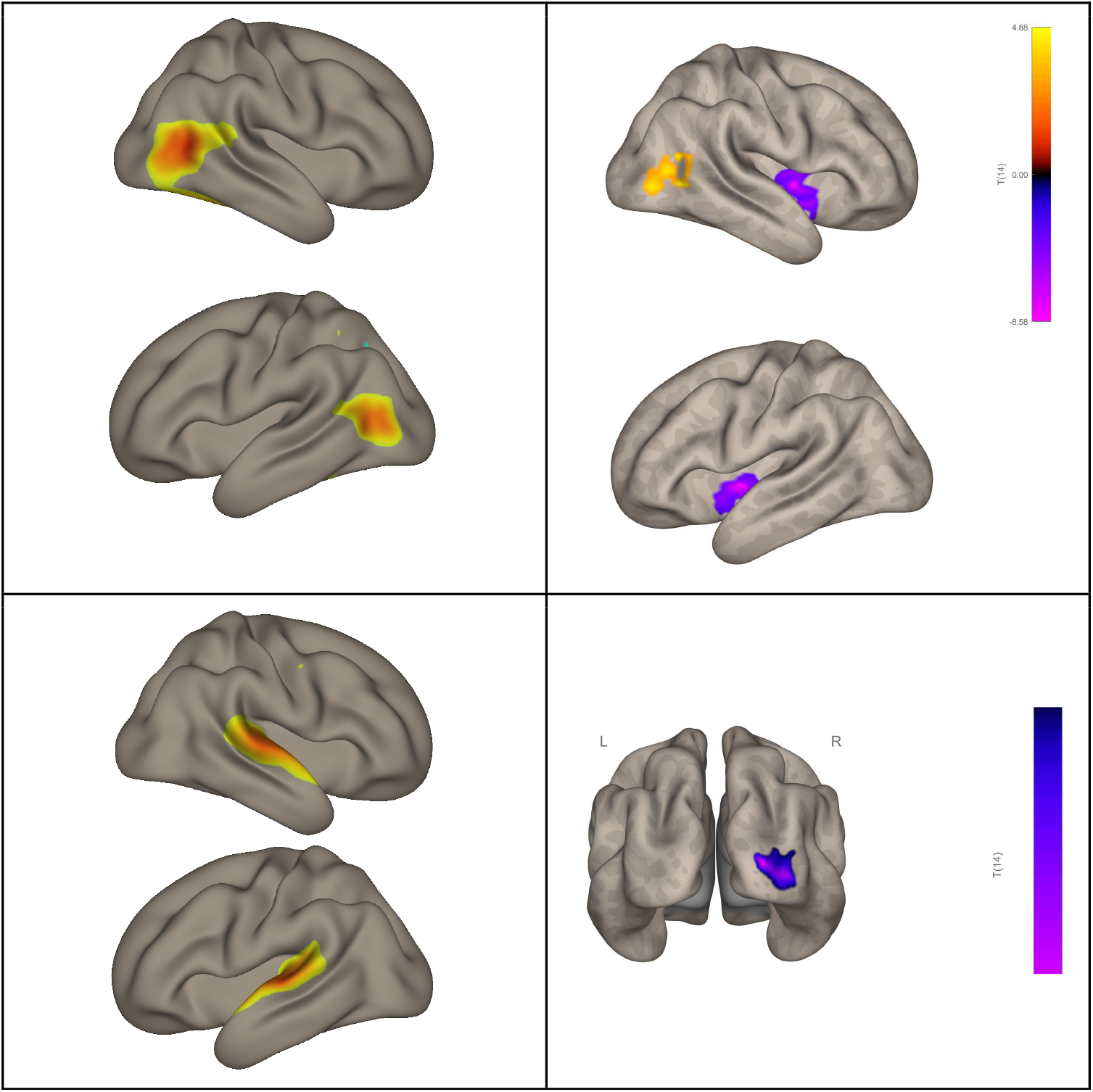
Results of the functional connectivity analysis of rs-fMRI data. if both seed and target are the same color (yellow-red or purple), this indicates increased FC; if they are different colors, this indicates decreased FC.

## Discussion

As reported in a series of our previous studies using an in-house set-up (Ciesla 2019, 2022, 2024), congruent low-frequency speech-extracted vibrations on fingertips can improve speech perception in difficult acoustic conditions, and especially after short audio-tactile training. Here, our resting-state fMRI findings provide insight into the brain networks likely engaged in the new experience of audio-tactile speech perception and those promoting training-related improvements in comprehension.

The revealed functional connectivity (FC) results show several effects when pre- and post-training resting-state networks are compared, which can be summarized as follows. First, we show a switch in functional connectivity of the lateral occipital cortex (LOC) from increased FC with bilateral dorsal anterior insula, to increased internal connectivity (i.e. of the right LOC with itself). Second, there is a change in functional connectivity of the early visual system, from one with the auditory system before training, to one with a hub in the left association somatosensory/multisensory cortex afterwards. And third, the sensorimotor system in the right hemisphere shows increased internal FC after training. While the effects involving the auditory and the sensorimotor system were somewhat expected following an audio-tactile intervention, the engagement of the visual system was less so initially.

We speculate that the revealed “visual” regions (both early V1-V2 and LOC) might represent parts of a network typically reported for *audio-visual* speech comprehension (e.g. Hertrich et al. 2020; Beer et al. 2013; Nath et al. 2012, Hauswald et al. 2018). This indicates that the brain functional system engaged in the utterly novel task of processing *audio-tactile* speech might be first built upon an existing blueprint for connectivity. In the current experiment, the speech task required comprehension of sentences in a non-native language that were also vocoded (resembling cochlear implant stimulation), and presented against background speech-noise, all rendering the task rather challenging. In everyday situations, when exposed to ambiguous speech, either itself distorted or in a challenging acoustic environment, we naturally search for informative visual cues, such as from lip-reading. These audio-visual associations develop from early years of life and are practiced on a regular basis throughout lifetime (Hickok et al. 2018, Doperiala et al. 2023). Beyond that, involvement of early visual cortex has been reported before for auditory and tactile (e.g. Braille reading) language tasks as well. Several studies have demonstrated this effect in both blind and sighted individuals, indicating that early visual regions have the capacity for amodal language processing (Amedi et al. 2003; Bedny et al. 2015; Reich et al. 2011, Seydell-Greenwald et al. 2023; Merabet et al. 2008). Here, after audio-tactile speech training, once it becomes apparent that no informative visual cues are available during task performance, the pre-existing (and potentially automatic in the context of speech perception) connectivity between the auditory system and the early visual system decreases (last row in Figure 2).

The other identified “visual” hub in the lateral occipital/temporal lobe encompassed regions such as the middle temporal visual area (MT), Extrastriate Body Area (EBA) and LOC. These brain areas, among other functions, can play a role in non-verbal communication, and specifically in encoding lip movements, body gestures, facial expressions and other language-related spatial cues (Kitada et al. 2014; Hickok et al. 2018, Van der Stoep et al. 2020). All these cues can facilitate comprehension in difficult acoustic conditions. Furthermore, the revealed adjacent pSTS is recognized as a key hub for multisensory integration, and particularly of audio-tactile and audio-visual inputs. It specifically specializes in processing temporally changing signals, such as, e.g. speech (King et al. 2019, Hertrich et al. 2020; Beer et al. 2013, Beauchamp et al. 2004).

In a non-speech specific context, LOC has also been shown to have multisensory properties (although mainly audio-visual), and is implicated in activities such as grasping and object recognition (e.g., Kassuba et al., 2013; Jao et al., 2015; Brand et al., 2020), as well as in response to passive vibrotactile stimulation on the hand (potentially related to the mechanism of object manipulation through increased connectivity with SI; Tal et al., 2016). Indeed, here the task might be considered as involving both language aspects, as well as body awareness and “reaching behavior”, as the goal is to combine sources of information arriving both from the ears and the fingertips inserted in the tactile device on the side of the body at the same time.

LOC is also shown after training, but less so in communication with bilateral anterior insular cortex. This outcome may stem from the insula’s role in various cognitive processes, particularly in the context of developing new skills or adaptation to new environments (e.g. Gogolla et al. 2017, Zuhlsdorf et al. 2023, Ardilla et al. 2014). Specifically, the revealed dorsal anterior part of insula, through its multiple connections with the rest of the brain, has been found to participate in language processing and general learning (e.g., Ardilla et al. 2014, although see: Woolnough et al. 2019), cognitive control, spatial attention tasks, as well as multimodal integration (as opposed to the social-emotional functions mostly attributed to the posterior and ventral-anterior aspects of the insula; Kurth et al. 2010; Uddin et al. 2018). In addition, the anterior insula may play a role in bodily-ownership, bodily self-awareness, and tactile perception. One suggested mechanism is through direct projections from the secondary somatosensory cortex to the insular cortex, which in turn innervates regions of the temporal lobe believed to be critical for tactile learning and memory (Kurth et al. 2010; Augustine [ed.], 2023). The decreased involvement of insula possibly reflects the progress in learning to use the newly available tactile inputs for improved speech comprehension.

Considering the role of the sensorimotor system, we demonstrate two additional effects. First, the applied audio-tactile speech training results in increased connectivity from the bilateral early visual cortex, to a posterior parietal region on the side opposite to the vibrotactile stimulation. Part of the revealed hub can be considered the association somatosensory cortex, where the experienced low-frequency tactile signal would be examined for its temporal and speech features. Other tactile operations attributed to this area include Braille reading, tactile attention, shape recognition, etc. (e.g. Kim et al 2014, Bodegard et al., 2001; Li Hegner et al., 2010). At the same time, the superior parietal lobe (constituting 40% of the revealed cluster) receives inputs not only from the early sensory system, and particularly the hand, but also from other modalities, which are further combined to inform behaviors, such as reaching or grasping (Sheth et al. 2014; Goodale and Milner, 1992). Finally, the demonstrated left-hemisphere SMG plays a critical role in the phonological analysis of language inputs, and also has multisensory properties (eg. Oberhuber et al. 2016). The increased direct functional connectivity after training between the early visual and the sensorimotor system/multisensory is not a straightforward result. It is possible that it reflects a mechanism similar to the one reported in some other works during tactile perception, though with opposite directions (Tal et al. 2016, Merabet et al. 2007). Specifically, Tal and colleagues showed deactivation of early visual regions (e.g. V1-V2) when the participants experienced passive delicate brushing of various body parts (although see a single-case EcoG study by Gaglianese and colleagues [2020] showing positive involvement of V1 when brushed), while Merebet and colleagues found active engagement of V1 in estimating roughness or inter-dot spacing in tactile patterns. This indicates that the responsiveness of early visual regions during tactile stimulation might be task-specific, and possibly reflect a certain mechanism of either motion processing or spatial analysis/tactile-to-visual remapping of inputs. In line with other authors, the increased FC with the early visual areas after training and its general role in the current experiment might be also mediated by mental imagery or visualization of the vibrating devices (eg. Reiner et al. 2008).

In addition after training we show increased internal connectivity within a right-hemisphere sensorimotor hub (ipsilateral to the tactile finger stimulation). The revealed region encompasses the primary (SI), secondary (SII) and association somatosensory cortices, including areas representing fingers, as well as the corresponding motor cortex. This finding can be related to several mechanisms. First, even though the responses to tactile stimuli are found to be mainly contralateral (and especially in SI), both animal and human neuroimaging studies show modulation of responses in ipsilateral SII (and to a lesser extent also SI) as well, possibly through dense transcallosal connectivity. Moreover, bilateral association sensorimotor cortex in the posterior parietal cortex is the area where inputs from both hands become integrated (Pala & Stanley 2022; Tame et al 2012, Hlushchuk et al. 2006). The sensorimotor cortex is also densely connected with the motor cortex, and specifically the regions representing hands, which enables successful manipulation of objects. The sensori-motor interactions involve the motor cortex responding to tactile stimulation, as well as the sensory cortex receiving feedback from and regulating motor behavior (Mastria et al. 2023, Tame et al. 2015, Bao et al. 2024). Several studies have demonstrated ipsilateral inhibition in the sensorimotor system within areas corresponding to specific body parts following unilateral tactile stimulation and during coordinated bilateral hand actions. This mechanism is believed to enhance the relevant sensory pathway by improving differentiation between left and right tactile or motor experiences (Hlushchuk et al. 2006, Tame et al. 2012, Zanona et al. 2022). It is possible that the effect observed here stems from this behaviourally relevant mechanism.

In summary, the main results of the study point to the role of the visual network, the auditory system, the sensorimotor system, and the insula, in audio-tactile integration of speech information and learning using our specific multisensory set-up. The findings are in partial agreement with our hypotheses (e.g. by showing engagement of the early sensory systems, as well as the pSTS and SMG in multisensory integration), even though no strengthening of direct connectivity was found between the auditory and the sensory cortex. The involvement of the early visual system, potentially also in the auditory-to-tactile communication (and thus integration of inputs) indicates that plastic changes following perceptual learning might be constrained by the existing connections established through experience (Makin & Krakauer 2023, Heimler & Amedi, 2020). At the same time, some of the revealed results in the visual, sensorimotor and insular networks can be interpreted in the framework of bodily awareness, grasping, and mental imagery, as the task required successful integration of inputs arriving from different spatial locations. Since the audio-tactile speech context is novel, never experienced in lifetime or in evolution, as opposed to the well-trained audio-visual speech context, a longer training regime might be necessary to modify the pre-existing connectivity patterns.

## Acknowledgements

This research was supported by Polish Ministry of Science and Higher Education MOBILNOŚĆ PLUS V grant (1642/MOB/V/2017/0) to K.C., ERC Consolidator Grant (773121 NovelExperiSense) to A.A., Horizon GuestXR (101017884) grant to AA; Israeli Science Foundation grant (3709/24) to A.A.

## References

1. Amedi A, Raz N, Pianka P, Malach R, Zohary E. Early ‘visual’ cortex activation correlates with superior verbal memory performance in the blind. Nat Neurosci. 2003 Jul;6(7):758–66.

2. Ardilla A, Bernal B, Rosselli M. Participation of the insula in language revisited: A meta-analytic connectivity study. Journal of Neurolinguistics, 29, 31–41; 2019.

3. Bamidis PD, Vivas AB, Styliadis C, Frantzidis C, Klados M, Schlee W, Siountas A, Papageorgiou SG. A review of physical and cognitive interventions in aging. Neurosci Biobehav Rev. 2014 Jul;44:206–20.

4. Bao S, Wang Y, Escalante YR, Li Y, Lei Y. Modulation of Motor Cortical Inhibition and Facilitation by Touch Sensation from the Glabrous Skin of the Human Hand. eNeuro.

5. Beauchamp MS, Lee KE, Argall BD, Martin A. Integration of auditory and visual information about objects in superior temporal sulcus. Neuron. 2004;41:809– 823.

6. Bedny M, Richardson H, Saxe R. “Visual” Cortex Responds to Spoken Language in Blind Children. J Neurosci. 2015 Aug 19;35(33):11674–81.

7. Beer A. L., Plank T., Meyer G., Greenlee M. W. (2013). Combined diffusion-weighted and functional magnetic resonance imaging reveals a temporal-occipital network involved in auditory-visual object processing. Front. Integr. Neurosci. 7:5.

8. Bernstein LE, Auer ET Jr, Eberhardt SP, Jiang J. Auditory Perceptual Learning for Speech Perception Can be Enhanced by Audiovisual Training. Front Neurosci. 2013;7:34

9. Bernstein LE, Auer ET Jr, Eberhardt SP, Jiang J. Auditory Perceptual Learning for Speech Perception Can be Enhanced by Audiovisual Training. Front Neurosci. 2013 Mar 18;7:34.

10. Biswal, B., Yetkin, F. Z., Haughton, V. M., and Hyde, J. S. (1995). Functional connectivity in the motor cortex of resting human brain using echo-planar MRI. Magn. Reson. Med. 34, 537–541.

11. Bodegard, A., Geyer, S., Grefkes, C., Zilles, K., and Roland, P. E. (2001). Hierarchical processing of tactile shape in the human brain. Neuron 31, 317–328.

12. Bolanowski Jr, S. J., Gescheider, G. A., Verrillo, R. T., & Checkosky, C. M. (1988). Four channels mediate the mechanical aspects of touch. The Journal of the Acoustical society of America, 84(5), 1680–1694.

13. Branco P, Seixas D, Deprez S, Kovacs S, Peeters R, Castro SL, Sunaert S. Resting-State Functional Magnetic Resonance Imaging for Language Preoperative Planning. Front Hum Neurosci. 2016 Feb 1;10:11.

14. Brand J, Piccirelli M, Hepp-Reymond MC, Eng K, Michels L. Brain Activation During Visually Guided Finger Movements. Front Hum Neurosci. 2020 Aug 14;14:309.

15. Cao, W., Luo, C., Zhu, B., Zhang, D., Dong, L., Gong, J., et al. (2014). Resting-state functional connectivity in anterior cingulate cortex in normal aging. Front. Aging Neurosci. 6:280.

16. Casserly E.D., Pisoni D.B. (2015). Auditory Learning Using a Portable Real-Time Vocoder: Preliminary Findings. Journal of Speech, Language and Hearing Research, 58(3), 1001–1016.

17. Cieśla, K., Wolak, T., Lorens, A., Heimler, B., Skarżyński, H., & Amedi, A. (2019). Immediate improvement of speech-in-noise perception through multisensory stimulation via an auditory to tactile sensory substitution. Restorative neurology and neuroscience, 37(2), 155–166.

18. Cieśla, K., Wolak, T., Lorens, A., Mentzel, M., Skarżyński, H., & Amedi, A. (2022). Effects of training and using an audio-tactile sensory substitution device on speech-in-noise understanding. Scientific Reports, 12(1), 1–16.

19. Cieśla, K. Wolak, T. Amedi, A. Neuronal basis of audio-tactile speech perception (2024). BIORXIV/2024/608369

20. Dopierała, A.A.W., Pérez, D.L., Mercure, E. et al. The Development of Cortical Responses to the Integration of Audiovisual Speech in Infancy. Brain Topogr, 36, 459–475 (2023).

21. Erb J., Molly H., Eisner F., Obleser J. (2013). The Brain Dynamics of Rapid Perceptual Adaptation to Adverse Listening Conditions. Journal of Neuroscience, 33(26), 10688–10697.

22. Fletcher, M. D., Hadeedi, A., Goehring, T., & Mills, S. R. (2019). Electro-haptic enhancement of speech-in-noise performance in cochlear implant users. Scientific Reports, 9(1), 1–8.

23. Fox MD, Greicius M. Clinical applications of resting state functional connectivity. Front. Syst. Neurosci. 4:19.

24. Gaglianese, A., Branco, M.P., Groen, I. et al. Electrocorticography Evidence of Tactile Responses in Visual Cortices. Brain Topogr. 33, 559–570 (2020).

25. Galvin K.L., Cowan R. S. C., Sarant J.Z., Alcantara J., Blarney P.J., Clark G.M. (1991). Use of a Multichannel Electrotactile Speech Processor by Profoundly Hearing-Impaired Children in a Total Communication Environment. Journal of the American Academy of Audiology, 12, 214–225.

26. George J. Augustine, et al. [eds]. Neuroscience (Sinauer Associates, Oxford University Press; 7th edition (March 1, 2023).

27. Gogolla N. The insular cortex. Curr Biol. 2017;27(12):R580–R586.

28. Goodale MA, Milner AD. Separate visual pathways for perception and action. Trends Neurosci. 1992 Jan;15(1):20–5.

29. Gori, M., Vercillo, T., Sandini, G., & Burr, D. (2014). Tactile feedback improves auditory spatial localization. Frontiers in Psychology, 5, 109563.

30. Guediche S, Blumstein SE, Fiez JA, Holt LL. Speech perception under adverse conditions: insights from behavioral, computational, and neuroscience research. Front Syst Neurosci. 2014 Jan 3;7:126.

31. Heimler B, Amedi A. Are critical periods reversible in the adult brain? Insights on cortical specializations based on sensory deprivation studies. Neurosci Biobehav Rev. 2020 Sep;116:494–507.

32. Hertrich, I., Dietrich, S., & Ackermann, H. (2020). The Margins of the Language Network in the Brain. Frontiers in Communication.

33. Hickok G, Rogalsky C, Matchin W, Basilakos A, Cai J, Pillay S, Ferrill M, Mickelsen S, Anderson SW, Love T, Binder J, Fridriksson J. Neural networks supporting audiovisual integration for speech: A large-scale lesion study. Cortex. 2018.

34. Hlushchuk Y, Hari R. Transient suppression of ipsilateral primary somatosensory cortex during tactile finger stimulation. J Neurosci. 2006 May 24;26(21):5819–24.

35. Huang J., Sheffield B., Lin. P., Zeng F.G. (2017). Electro-Tactile Stimulation Enhances Cochlear Implant Speech Recognition in Noise. Science Reports, 7(1), 2196.

36. Jao RJ, James TW, James KH. (2015) Crossmodal enhancement in the LOC for visuohaptic object recognition over development. Neuropsychologia. 2015 Oct;77:76–89.

37. Kassuba T. Menz M.M. Röder B. Siebner H.R. Multisensory interactions between auditory and haptic object recognition.Cereb. Cortex. 2013; 23:1097–1107.

38. Katsuno Y, Ueki Y, Ito K, Murakami S, Aoyama K, Oishi N, Kan H, Matsukawa N, Nagao K, Tatsumi H. Effects of a new speech support application on intensive speech therapy and changes in functional brain connectivity in patients with post-stroke aphasia. Front Hum Neurosci. 2022 Sep 22;16:870733.

39. Kim J, Müller KR, Chung YG, Chung SC, Park JY, Bülthoff HH, Kim SP. Distributed functions of detection and discrimination of vibrotactile stimuli in the hierarchical human somatosensory system. Front Hum Neurosci. 2015 Jan 21;8:1070.

40. King, A.J., Hammond-Kenny, A., Nodal, F.R. (2019). Multisensory Processing in the Auditory Cortex. In: Lee, A., Wallace, M., Coffin, A., Popper, A., Fay, R. (eds) Multisensory Processes. Springer Handbook of Auditory Research, vol 68. Springer, Cham.

41. Kitada R, Yoshihara K, Sasaki AT, Hashiguchi M, Kochiyama T, Sadato N. The brain network underlying the recognition of hand gestures in the blind: the supramodal role of the extrastriate body area. J Neurosci. 2014 Jul 23;34(30):10096–108.

42. Landelle C, Caron-Guyon J, Nazarian B, Anton JL, Sein J, Pruvost L, Amberg M, Giraud F, Félician O, Danna J, Kavounoudias A. Beyond sense-specific processing: decoding texture in the brain from touch and sonified movement. iScience. 2023 Sep 20;26(10):107965.

43. Lemée JM, Berro DH, Bernard F, Chinier E, Leiber LM, Menei P, Ter Minassian A. Resting-state functional magnetic resonance imaging versus task-based activity for language mapping and correlation with perioperative cortical mapping. Brain Behav. 2019 Oct;9(10):e01362. doi: 10.1002/brb3.1362.

44. Li Hegner, Y., Lee, Y., Grodd, W., and Braun, C. (2010). Comparing tactile pattern and vibrotactile frequency discrimination: a human FMRI study. J. Neurophysiol. 103, 3115–3122.

45. Liu C, Jiao L, Li Z, Timmer K, Wang R. Language control network adapts to second language learning: A longitudinal rs-fMRI study. Neuropsychologia. 2021 Jan 8;150:107688

46. Makin TR, Krakauer JW (2023) Against cortical reorganization eLife 12:e84716.

47. Maruyama T, Takeuchi H, Taki Y, Motoki K, Jeong H, Kotozaki Y, Nakagawa S, Nouchi R, Iizuka K, Yokoyama R, Yamamoto Y, Hanawa S, Araki T, Sakaki K, Sasaki Y, Magistro D, Kawashima R. Effects of Time-Compressed Speech Training on Multiple Functional and Structural Neural Mechanisms Involving the Left Superior Temporal Gyrus. Neural Plast. 2018 Feb 20;2018:6574178.

48. Mastria G, Scaliti E, Mehring C, Burdet E, Becchio C, Serino A, Akselrod M. Morphology, Connectivity, and Encoding Features of Tactile and Motor Representations of the Fingers in the Human Precentral and Postcentral Gyrus. J Neurosci. 2023 Mar 1;43(9):1572–1589.

49. Merabet LB, Swisher JD, McMains SA, Halko MA, Amedi A, Pascual-Leone A, Somers DC. Combined activation and deactivation of visual cortex during tactile sensory processing. J Neurophysiol. 2007 Feb;97(2):1633–41.

50. Merabet LB, Hamilton R, Schlaug G, Swisher JD, Kiriakopoulos ET, Pitskel NB, Kauffman T, Pascual-Leone A. Rapid and reversible recruitment of early visual cortex for touch. PLoS One. 2008 Aug 27;3(8):e3046.

51. Nath AR, Beauchamp MS. A neural basis for interindividual differences in the McGurk effect, a multisensory speech illusion. Neuroimage. 2012 Jan 2;59(1):781–7.

52. Ngo GH, Khosla M, Jamison K, Kuceyeski A, Sabuncu MR. Predicting individual task contrasts from resting-state functional connectivity using a surface-based convolutional network. Neuroimage. 2022 Mar;248:118849.

53. Nilsson M., Soli S.D., Sullivan J.A. (1994). Development of the Hearing In Noise Test for the measurement of speech reception thresholds in quiet and in noise. Journal of the Acoustical Society of America, 95(2), 1085–1099.

54. O’Sullivan AE, Crosse MJ, Liberto GMD, de Cheveigné A, Lalor EC. Neurophysiological Indices of Audiovisual Speech Processing Reveal a Hierarchy of Multisensory Integration Effects. J Neurosci. 2021;41(23):4991–5003.

55. Oberhuber M, Hope TMH, Seghier ML, Parker Jones O, Prejawa S, Green DW, Price CJ. Four Functionally Distinct Regions in the Left Supramarginal Gyrus Support Word Processing. Cereb Cortex. 2016 Oct 1;26(11):4212–4226.

56. Oberhuber M, Hope TMH, Seghier ML, Parker Jones O, Prejawa S, Green DW, Price CJ. Four Functionally Distinct Regions in the Left Supramarginal Gyrus Support Word Processing. Cereb Cortex. 2016 Oct 1;26(11):4212–4226.

57. Occelli, V., Spence, C., & Zampini, M. (2009). Compatibility effects between sound frequency and tactile elevation. Neuroreport, 20(8), 793–797.

58. Pala A, Stanley GB. Ipsilateral Stimulus Encoding in Primary and Secondary Somatosensory Cortex of Awake Mice. J Neurosci. 2022 Mar 30;42(13):2701–2715.

59. Pisciottano, F., Cinalli, A. R., Stopiello, J. M., Castagna, V. C., Elgoyhen, A. B., Rubinstein, M., (2019). Inner ear genes underwent positive selection and adaptation. Molecular Biology and Evolution, 36(8), 1653–1670.

60. Price CJ. A review and synthesis of the first 20 years of PET and fMRI studies of heard speech, spoken language and reading. Neuroimage. 2012 Aug 15;62(2):816–47.

61. Răutu, I.S., De Tiège, X., Jousmäki, V. et al. Speech-derived haptic stimulation enhances speech recognition in a multi-talker background. Sci Rep 13, 16621 (2023).

62. Raichle ME. A paradigm shift in functional brain imaging. J Neurosci. 2009;29:12729–12734.

63. Reich L, Szwed M, Cohen L, Amedi A. A ventral visual stream reading center independent of visual experience. Curr Biol. 2011 Mar 8;21(5):363–8.

64. Reiner M. Seeing through touch: the role of haptic information in visualization. In: Gilbert J.K., Reiner M., Nakhleh M.B., editors. Visualization: Theory and Practice in Science Education. Springer; Netherlands: 2008. pp. 73–84.

65. Renier LA, Anurova I, De Volder AG, Carlson S, VanMeter J, Rauschecker JP. Multisensory integration of sounds and vibrotactile stimuli in processing streams for “what” and “where”, J Neurosci, 2009, vol. 29, 10950–10960.

66. Robertson FC, Roos A, Douglas TS, Stein DJ, Meintjes EM. Ipsilateral functional deactivation to unilateral sensorimotor tasks measured with fMRI and NIRS/DOT; OHBM 2014.

67. Russo, F. A., Ammirante, P., & Fels, D. I. (2012). Vibrotactile discrimination of musical timbre. Journal of Experimental Psychology: Human Perception and Performance, 38(4), 822.

68. Schnupp, J., Nelken, I., & King, A. (2011). Auditory neuroscience: Making sense of sound. MIT press.

69. Schulte A, Marozeau J, Ruhe A, Büchner A, Kral A, Innes-Brown H. Improved speech intelligibility in the presence of congruent vibrotactile speech input. Sci Rep. 2023;13(1):22657.

70. Seydell-Greenwald, A, Wang X, Newport EL, Bi Y, Striem-Amit E. Spoken language processing activates the primary visual cortex. PLoS One. 2023 Aug 11;18(8):e0289671.

71. Sharp, A., Houde, M. S., Maheu, M., Ibrahim, I., and Champoux, F. (2019). Improve tactile frequency discrimination in musicians. Exp. Brain Res. 237, 1–6.

72. Sheth V, Young R. Ventral and dorsal streams in cortex: focal vs. ambient processing/exploitation vs. exploration Journal of Vision August 2014, Vol.14, 51.

73. Snir A, Cieśla K, Ozdemir G, Vekslar R, Amedi A. Localizing 3D motion through the fingertips: Following in the footsteps of elephants. iScience. 2024 Apr 26;27(6):109820.

74. Sun X, Li L, Ding G, Wang R, Li P. Effects of language proficiency on cognitive control: Evidence from resting-state functional connectivity. Neuropsychologia. 2019 Jun;129:263–275.

75. Tal, Z., Geva, R., & Amedi, A. (2016, February). The origins of metamodality in visual object area LO: Bodily topographical biases and increased functional connectivity to S1. NeuroImage, 127, 363–375.

76. Tamè L, Braun C, Lingnau A, Schwarzbach J, Demarchi G, Li Hegner Y, Farnè A, Pavani F. The contribution of primary and secondary somatosensory cortices to the representation of body parts and body sides: an fMRI adaptation study. J Cogn Neurosci. 2012 Dec;24(12):2306–20.

77. Tamè L, Pavani F, Braun C, Salemme R, Farnè A, Reilly KT. Somatotopy and temporal dynamics of sensorimotor interactions: evidence from double afferent inhibition. Eur J Neurosci. 2015 May;41(11):1459–65.

78. Tie, Y., Rigolo, L., Norton, I. H., Huang, R. Y., Wu, W., Orringer, O., et al. (2014). Defining language networks from resting-state fMRI for surgical planning – a feasibility study. Hum. Brain Mapp. 35, 1018–1030.

79. Uddin LQ, Nomi JS, Hébert-Seropian B, Ghaziri J, Boucher O. Structure and Function of the Human Insula. J Clin Neurophysiol. 2017 Jul;34(4):300–306.

80. Van Békésy G. (1959). Similarities between hearing and skin sensations. Psychological Review, 1–22.

81. Van der Stoep N, Alais D. Motion Perception: Auditory Motion Encoded in a Visual Motion Area. Curr Biol. 2020 Jul 6;30(13):R775–R778.

82. Walkowiak A., Kostek B., Lorens A., Obrycka A., Wasowski A., Skarzynski H. (2010). Spread of Excitation (SoE) - a non-invasive assessment of cochlear implant electrode placement. Cochlear Implants International, 11(1), 479–481.

83. Wang X, Wu Q, Egan L, Gu X, Liu P, Gu H, Yang Y, Luo J, Wu Y, Gao Z, Fan J. Anterior insular cortex plays a critical role in interoceptive attention. Elife. 2019 Apr 15;8:e42265.

84. Weisenberger J.M., Percy M.E. (1995). The Transmission of Phoneme-Level Information by Multichannel Tactile Speech Perception Aids. Ear and Hearing, 16(4), 392–406.

85. Woolnough O, Forseth KJ, Rollo PS, Tandon N. Uncovering the functional anatomy of the human insula during speech. Elife. 2019 Dec 19;8:e53086.

86. de Freitas Zanona A, Romeiro da Silva AC, Baltar do Rego Maciel A, Shirahige Gomes do Nascimento L, Bezerra da Silva A, Piscitelli D, Monte-Silva K. Sensory and motor cortical excitability changes induced by rTMS and sensory stimulation in stroke: A randomized clinical trial. Front Neurosci. 2023, 16:985754.

87. Zhu, L., Fan, Y., Zou, Q., Wang, J., Gao, J. H., and Niu, Z. (2014). Temporal reliability and lateralization of the resting-state language network. PLoS ONE 9:e85880.

88. Zühlsdorff K, Dalley JW, Robbins TW, Morein-Zamir S. Cognitive flexibility: neurobehavioral correlates of changing one’s mind. Cereb Cortex. 2023 Apr 25;33(9):5436–5446.

